# Differentiation of airway cholinergic neurons from human pluripotent stem cells for airway neurobiology studies

**DOI:** 10.1101/2022.03.23.485449

**Authors:** P.A. Goldsteen, A.M. Sabogal Guaqueta, I.S.T. Bos, L.E.M. Kistemaker, L. van der Koog, M. Eggens, A.J. Halayko, A.M. Dolga, R. Gosens

## Abstract

Airway cholinergic nerves play a key role in airway physiology and disease. In asthma and other diseases of the respiratory tract, airway cholinergic neurons undergo plasticity and contribute to airway hyperresponsiveness and mucus secretion. We currently lack mechanistic understanding of airway cholinergic neuroplasticity due to the absence of human *in vitro* models. Here, we developed the first human *in vitro* model for airway cholinergic neurons using human pluripotent stem cell (hPSC) technology. hPSCs were differentiated towards mature and functional airway cholinergic neurons via a vagal precursor. Airway cholinergic neurons were characterized by ChAT and VAChT expression, and responded to chemical stimulation with changes in Ca^2+^ mobilization. Co-culture of hPSC-derived airway cholinergic neurons with airway smooth muscle cells enhanced phenotypic and functional characteristics of these neurons. The differentiation protocol we developed for human airway cholinergic neurons from hPSCs allows for studies into airway neurobiology and airway neuroplasticity in disease.

## Introduction

The lungs are innervated through a dense network of afferent and efferent nerves, which are arranged along the vagus nerve^1^. Among the efferent nerves, the parasympathetic neurons are most dominant in controlling airway smooth muscle (ASM) tone. Parasympathetic cholinergic nerves are major regulators of bronchoconstriction and mucus secretion^2^. Like all neurons, the airway nervous system is subjected to changes over time in response to intrinsic and extrinsic stimuli, known as neuronal plasticty^3,4^. Severe or prolonged stimuli can cause permanent changes to the neurons, manifested as altered neurite length or innervation, lowered firing threshold, or even phenotype switching^5,6^. Asthma patients have an increased innervation of both the sensory and the autonomic cholinergic nervous system^7,8^. In asthma, neuroplasticity of the cholinergic nervous system is a newly discovered phenomenon^8^. However, we can only speculate about the underlying mechanisms.

Conventional models like animal models or patient biopsies can only provide limited information about mechanisms underlying neuroplasticity^9^. Patient biopsies provide much information on the final stages of neuronal remodeling. Animal models can be favored over biopsies for mechanistic studies as they take into account full physiological complexity, and the nervous system is well integrated into the organs and connected to the central nervous system (CNS). However, the problem with animal models is that the translation to the human situation is poor^10,11^. Human pluripotent stem cells (hPSCs) can aid in the development of a human disease model to study the process of neuronal plasticity *in vitro*^9^. Developing a human *in vitro* model to understand neuroplasticity in asthma has mostly been hampered by a lack of protocols for robust differentiation from hPSCs to that of airway neurons.

Airway cholinergic neurons and enteric neurons originate from a vagal neural crest cell (NCC) precursor before developing into different directions^12,13^. Innervation of the gastrointestinal tract by enteric neurons is better characterized than the respiratory innervation by intrinsic neurons^12^. The differentiation of NCCs towards the peripheral neurons is dependent on neurotrophic factor (NTF) signaling. NTFs regulate neurogenesis, neuronal differentiation, neuronal survival, nerve conduction, and neuronal plasticity^12,14^. Brain-derived neurotrophic factor (BDNF) is the predominant NTF in the lungs^15^, opposed to glial cell-derived neurotrophic factor (GDNF) in the enteric system^16,17^. Secreted NTFs act as essential chemo-attractants for NCCs: ASM secretes BDNF in the lungs and guides airway neurons^18^. Based on this information, we hypothesized that transforming existing protocols for enteric neuron differentiation into a protocol for airway neuronal differentiation is possible^9,17^.

In this study, we established a robust protocol for airway cholinergic neurons differentiation. Using dual SMAD inhibition and Wnt activation, p75^+^-HNK1^+^ vagal neural crest cell (NCC) precursors were generated, confirmed by HOXB3 and HOXB5 expression. Subsequently, vagal NCCs were guided into mature and functional airway cholinergic neurons using BDNF. Obtained neurons expressed VAChT and ChAT, as well as the cholinergic M3 receptor, and responded to acetylcholine (ACh) and potassium chloride (KCl) with changes in Ca^2+^ mobilization. In addition, co-culturing hPSC-derived airway cholinergic neurons with ASM cells showed neuronal innervation and demonstrated the possibility of culturing hPSC-derived neurons in close proximity to cells of the local microenvironment. Lastly, to implement the generated neurons into a disease model, hPSC-derived airway neurons were stimulated with dexamethasone or a cocktail of Th2 cytokines. Dexamethasone stimulation reduced the yield of obtained cholinergic neurons. Cytokine stimulation did not hamper the viability of the neurons. Developing such a human in *vitro* disease model will allow us to study airway neurobiology and neuronal plasticity, and propose therapeutic targets in lung diseases.

## Results

### Derivation of vagal NCCs from hPSCs

During embryonic development, airway cholinergic neurons originate from vagal NCCs. H9WA09 cells were directed into vagal NCCs using dual SMAD inhibition and early WNT activation using chemically defined conditions (Figure 1A)^17^. To induce dual SMAD inhibition, SB431542 and LDN193189 were used as a BMP/TGF-β pathway inhibitor and a BMP pathway inhibitor, respectively. Later during the NCC differentiation, CHIR99021 was added for temporal WNT activation. A monolayer of pluripotent H9WA09 colonies differentiated into vagal NCCs in 12 days (Figure 1B). Following vagal NCC induction, cells were cultured as spheroids for four days in order to further mature the NCC phenotype (Figure 1B, right panel). On day 0, colonies were positive for OCT4, a marker for pluripotency (Figure 1C), while after 16 days of differentiation, cells showed decreased OCT4 expression and shifted to SOX10^+^ NCCs (Figure 1D). The NCC stem cell marker Nestin was detected in day 16 spheroids (Supplementary Figure 1A).

**Figure 1.**
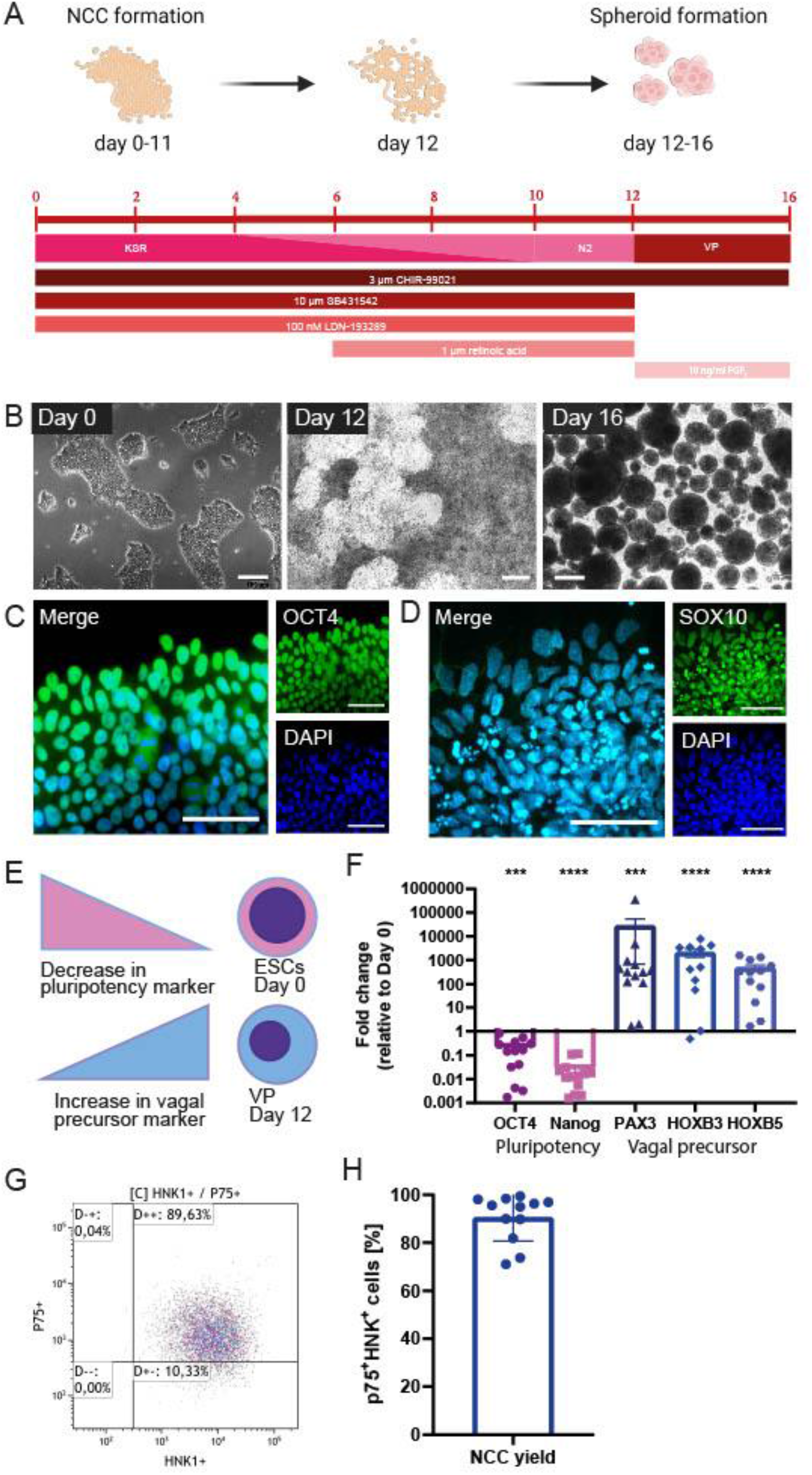
Derivation of vagal NCCs from hPSCs. A. Schematic overview of vagal NCC induction from hPSCs. H9WA09 cells were directed into a vagal NCC faith using dual SMAD inhibition, and early WNT activation, r.a. was added to direct NCCs from a cranial into a more distal phenotype. Knock-out serum replacer medium (KSR) was gradually replaced by N2 medium. Vagal NCCs were cultured in spheroids for four days to complete maturation. Created with BioRender.com. B. Bright images of the differentiating cells over time. Scale bar = 200 µm. C. OCT4 expression of Day 0 pluripotent cells. Scale bar = 50 µm. D. SOX10 expression of Day 16 vagal NCC spheroids. Scale bar = 50 µm. E-F. Gene expression of several markers for pluripotency, NCC, and vagal NCC. After 12 days, a decrease in the pluripotency genes OCT4 and Nanog was observed. The NCC marker PAX3 was upregulated, in combination with the vagal NCC markers HOXB3 and HOXB5. (N = 13) G-H. FACS analysis showed that 90.48% (± 2.83%) of the vagal NCCs were p75+-HNK1+ double positive. (N = 12) A paired t-test was performed to calculate statistical significance between means. ***p<0.001; ****p<0.0001 compared to day 0.

To confirm the vagal NCC identity of the acquired cells, we performed a gene expression analysis of NCC (*PAX3*) and vagal NCC (*HOXB3, HOXB5*) markers. Firstly, *PAX3* is reported to be one of the earliest markers of NCC induction^22^. In addition, vagal NCCs express *HOXB3* and *HOXB5* during embryogenesis^23,24^. After 12 days of differentiation, vagal NCCs displayed lower expression of pluripotency genes (*OCT4, p* < 0.001; *NANOG, p* < 0.0001) and higher expression of NCC genes (*PAX3, p* < 0.001; *HOXB3, p* < 0.0001; *HOXB5, p* < 0.0001) compared to day 0 (Figure 1E-F). On day 12, the normalized expression of OCT4 and NANOG was significantly downregulated compared to day 0 (Figure 1F). Comparison of these relative gene expressions between samples collected during differentiation revealed a population transition from pluripotent stem cells to vagal NCCs. *PAX3* is involved in NCC development; *PAX6*, on the other hand, is an important early marker for neural tube formation and subsequent central nervous system differentiation^22^. The obtained NCCs showed a high expression of *PAX3*, whereas *PAX6* was completely absent (Supplementary Figure 1B).

The efficiency of the differentiation protocol was analyzed by determining the yield of NCCs using FACS. HNK1 and p75 are both surface markers that are highly abundant on the surface of migratory NCCs^25^. Figure 1G shows a representative FACS analysis at day 12 of differentiation. Induction of H9WA09 cells towards a NCC fate was highly efficient, with 90.48% (± 2.83%) of total cells being positive for p75^+^HNK1^+^ (Figure 1H). Together, these data indicate that vagal NCCs (displaying HOXB3 and HOXB5) were induced with high efficiency.

### Neuronal differentiation from vagal NCCs

To differentiate vagal NCC precursors further into mature airway cholinergic neurons, we plated day-16-spheroids on FB/LM-coated surfaces (Figure 2A). A neuronal network was formed gradually and increased every day, with the first axonal outgrowth already visible after 24 hours (Figure 2B, left panel; Figure 2C). At day 25 of differentiation, a neuronal network was clearly visible, which further expanded and became denser over time. β-3-tubulin staining confirmed the overall neuronal network formation (Figure 2C-E).

**Figure 2.**
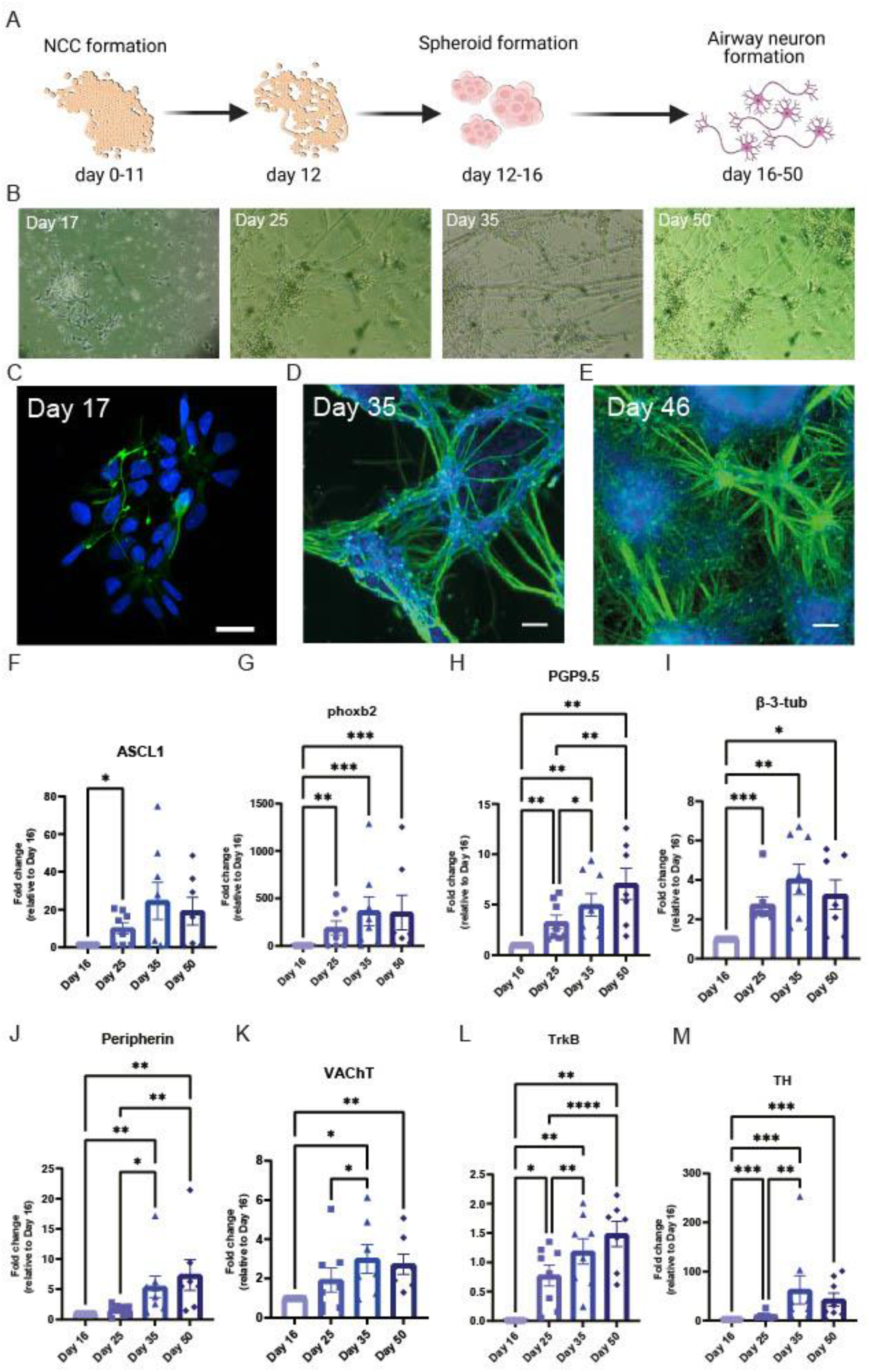
Differentiation of airway cholinergic neurons. A. Schematic overview of hPSC differentiation into airway cholinergic neurons. Created with BioRender.com. B. Brightfield images of the differentiating cells over time. C-E. Immunofluorescence images of differentiating cells over time, showing the development of β-3-tubulin expression. Over time a greater neuronal network is being formed. Day 17, scale bar = 20 µm. Day 35 and day 46, scale bar = 100 µm. F-L. Gene expression of airway cholinergic neuronal development over time. Gene expression was examined at different time points: Day 16, day 25, day 35, and day 50. ASCL1 (F) and phoxb2 (G) are essential for neuronal development and increased over the course of day 50. The pan-neuronal markers PGP9.5 (H) and β-3-tubulin (I) indicate the formation of neurons, in combination with peripherin (J) they indicate the development of peripheral neurons. VAChT (K), indicating the formation of cholinergic neurons, was higher expressed over the course of 50 days. TrkB (L) increased in expression over the course of 50 days. TH (M) showed a peak at day 35 before declining in expression towards day 50. (N = 8) A mixed-effect analysis followed by Tukey’s multiple comparisons test was performed to calculate statistical significance between means. *p<0.05; **p<0.01; ***p<0.001; ****p<0.0001, compared to day 16.

To track the development of airway neurons during differentiation, we compared the gene expression between samples collected at several time points during the differentiation protocol. *In utero*, vagal NCCs express PHOX2B and ASCL1 after their inclusion in the foregut ^26^. Accordingly, differentiated cells showed a relative increase in gene expression of PHOX2B, and ASCL1, with a peak at day 35 for ASCL1 (Figure 2F-G). Next, the expression of both the TUBB3 and UCHL1 showed that neurons were abundantly present and that their gene expression increased from day 16 to day 50 (Figure 2H-I). Expression of peripherin (*PRPH*), a cytoskeletal protein found in peripheral neurons, also increased towards day 50 and confirmed a peripheral neuronal phenotype (Figure 2J). Vesicular acetylcholine transporter (VAChT, *SLC18A3*) expression confirmed the cholinergic phenotype of the neurons (Figure 2K).

Similarly, Tropomyosin receptor kinase B (TrkB) is the predominant NTF receptor present in the airways and is therefore an excellent indicator of airway neurons identity. Both SLC18A3 and TrkB (*NTRK2*) increased over the course of 50 days, indicating neuronal formation and maturation (Figure 2K-L). TH is a marker for sympathetic neurons; however, it is also a precursor for cholinergic neurons during development^27^. Consistent with this contention, tyrosine hydroxylase (*TH*) peaked in expression at day 35 of differentiation and declined afterward (Figure 2M). Our data implies that the cells matured over time into peripheral cholinergic neurons as *TH* expression decreased (Figure 2M), and the expression of PRPH and SLC18A3 increased.

### BDNF induces cholinergic neuron differentiation

Further identification of neuronal identity revealed that after 50 days, most of the generated airway neurons were VAChT^+^ (Figure 3A), indicating a cholinergic phenotype. Neurons matured gradually into airway cholinergic neurons after 35 days of differentiation. On day 35, neurons stained positive for β-3-tubulin, while VAChT expression was sparse (Supplementary Figure 2A). Other important markers of the neuronal network include the presence of synaptophysin (SYP) and peripherin (Figure 3B-C). Most of the neurons appeared peripherin^+^ at day 50, indicating that the obtained neurons are PNS neurons, not CNS neurons (Figure 3C). Also, part of the neurons expressed SYP, indicating the presence of synapses. In autonomic neurons, synapses are distributed over the length of the axons, rather than having a synapse formed at the extremity end of the axons, which was also observed here (Figure 3B).

**Figure 3.**
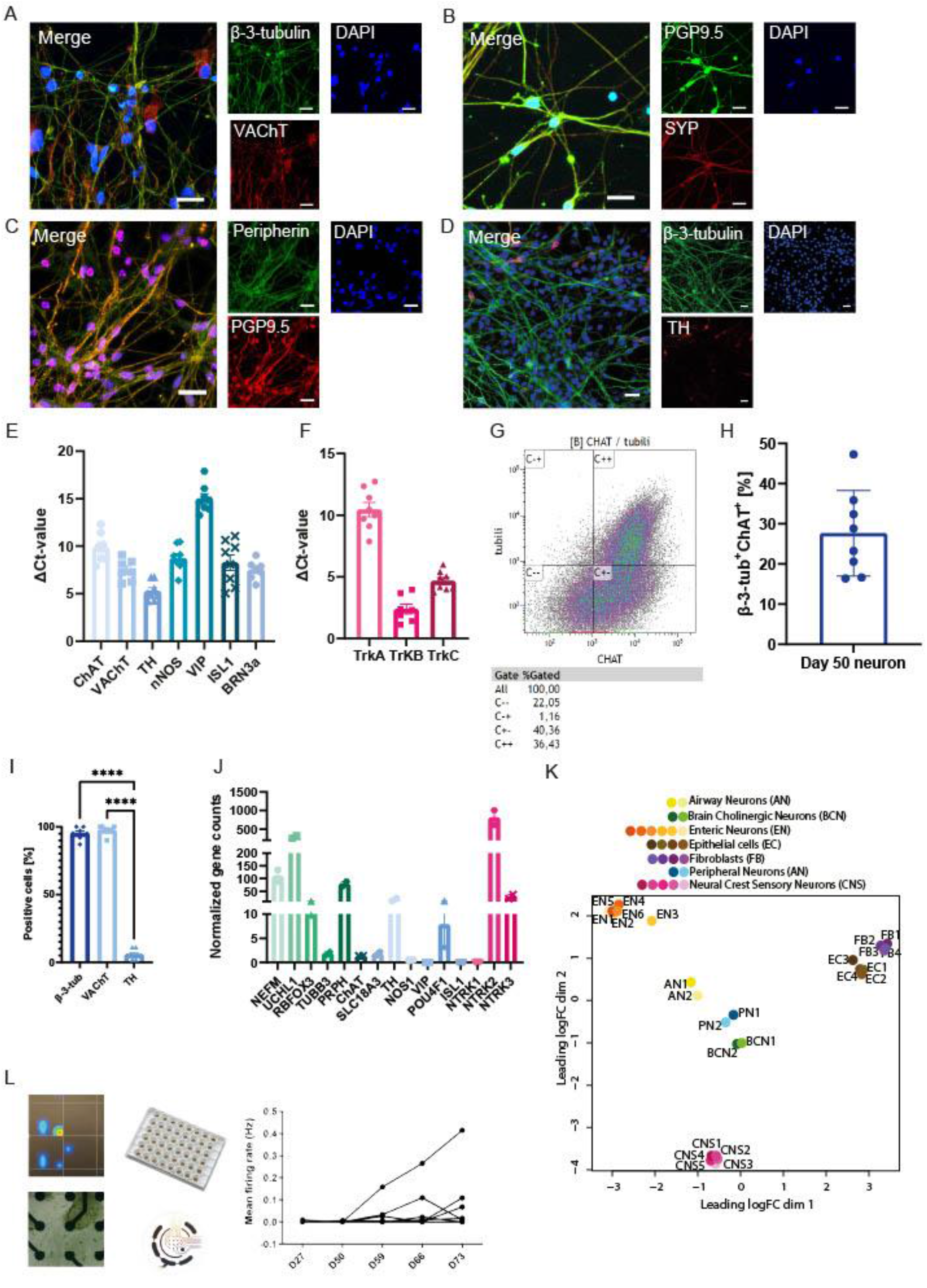
Airway cholinergic neurons. A-D. Immunofluorescence images characterizing neurons. β-3-tubulin or PGP9.5 were used as a pan-neuronal marker. The neurons show the presence of VAChT (A) after 50 days of differentiation, in addition to SYP (B) and peripherin (C). A smaller proportion of the neurons show TH^+^ after 50 days. E-F. Gene expression of neurons after 50 days of differentiation. The neurons show both ChAT and VAChT confirming their cholinergic identity (E), in addition to the presence of TH. A proportion of the neurons also expresses nNOS or the sensory markers ISL1 and BRN3a. VIP was hardly expressed in the neurons. TrkB was the predominant neurotrophic receptor present on the neurons. TrkC was also present, whereas TrkA was hardly expressed (N = 8). G-H. FACS analysis demonstrated that 27.7% (± 3.8%) of the neurons are β-3-tubulin^+^-ChAT^+^, indicative of cholinergic neurons. A representative FACS analysis plot is shown in G. (N = 8) I. Quantification of IF staining of β-3-tubulin, VAChT, and TH showed that the majority of cells expressed β-3-tubulin and VAChT, whilst TH was only sparsely present. J. RNA sequencing data of day 50 cholinergic neurons (N = 2). K. PCA analysis showed that hPSC-derived airway neurons were distant compared to airway epithelial cells or fibroblasts, but showed more overlap with hPSC-derived peripheral neurons or brain cholinergic neurons. L. MEA analysis of neurons. Neurites extend over the electrodes of the MEA plate, and started spontaneous firing after day 50 of differentiation and increased up to day 73. Figures were presented as mean ± SEM.) A one-way ANOVA followed by Tukey’s multiple comparisons test was performed to calculate statistical significance between means. ****p<0.0001.

Vagal NCCs migrate along with somites and are the precursor for various neuronal cell types during embryonic development, next to airway cholinergic neurons^13^. We analyzed the gene expression of autonomic neuronal markers like Choline acetyltransferase (ChAT, *CHAT*), *SLC18A3*, neuronal nitric oxide synthase (nNOS, *NOS1*), vasoactive intestinal peptide (*VIP*), and TH, as well as sensory neuronal markers like *ISL1* and BRN3a (*POU4F1*) to characterize the generated neuronal pool (Figure 3E). *TH* gene expression was most abundant, whereas *SLC18A3, CHAT, NOS1*, and sensory markers were similar in expression. Only *VIP* appeared to be absent in the generated neurons. As TH gene expression was abundant, we confirmed TH expression using IF staining. We found that only a minor proportion of the generated neurons was TH^+^ (Figure 3D and 3I), in contrast to abundant VAChT expression (Figure 3A and 3I). TH^+^ neurons were not overlapping with VAChT^+^ neurons (Supplementary Figure 2B). In addition, the TrkB receptor is the predominant receptor in the airways during development^18^. As expected, after 50 days of differentiation in the presence of BDNF, *NTRK2* was most expressed compared to *NTRK1* and *NTRK3* (Figure 3F).

Airway cholinergic neurons use ACh as their primary neurotransmitter, which would be confirmed by the presence of the ACh synthesizing enzyme ChAT. FACS analysis further confirmed the final cholinergic phenotype of day 50 of neuronal differentiation and maturation. After 50 days of differentiation, 27.6% (SEM ± 4.5%) of the generated neurons were β-3-tubulin^+^-ChAT^+^ (Figure 3G-H). When the proportion of β-3-tubulin^+^ cells was analyzed for ChAT expression, 80.0 % (SEM ± 6.5%) were ChAT^+^ cells. Quantification of the IF stainings of the generated neurons further confirmed that 95.1% (SEM ± 4.6%) of the cells were β-3-tubulin^+^ and 97.6% (SEM ± 3.3%) of the cells were VAChT^+^ (Figure 3I), supporting the conclusion that the majority of the neurons formed are cholinergic.

RNA sequencing analysis of two samples at day 50 of differentiation confirmed the RT-qPCR results. Neuronal markers were abundantly present and in higher numbers than other neural crest derivatives like glial cells, mesenchymal cells, melanocytes, or endocrine cells. The pan-neuronal markers *NEFM* and *UCHL1* showed the highest read counts (Figure 3J) compared to other cell types (Supplementary Figure 3A-C). Furthermore, the distribution of neuronal subtypes and Trk-receptors in the RNAseq data is similar to what we observed by PCR analysis, with a predominant expression of *TH* and *NTRK2*, respectively (Figure 3I). Furthermore, analysis of receptor expression provided more insight into the maturation of the obtained neurons. Mature airway cholinergic neurons display the autonomic ganglia-specific nicotinic and muscarinic ACh receptors: *CHRNA3, CHRNB4, CHRM2, CHRM3*, and *CHRM4*^28,29^. The ACh receptors, especially the nicotinic receptor 3 (*CHRNA3*) and muscarinic receptor 3 (*CHRM3*), appeared in the generated neurons. In addition, the glutamate receptor *VGLUT2* was abundantly expressed on day 50 neuronal differentiation, as well as the vesicular monoamine transporter *VMAT2I* (Supplementary Table 1).

We performed a comparison to determine the resemblance of the generated neurons with other cells types through principal component analysis (PCA). PCA is a statistical technique to summarize the information from extensive databases to improve interpretability while keeping as much information as possible^30^. We compared the RNAseq profile of the obtained neurons with previously published datasets using hPSC-derived peripheral neurons^31^, hPSC-derived neural crest sensory neurons^32^, human-induced brain cholinergic neurons (from fetal fibroblasts)^33^, human enteric neurons^34^, human epithelial cells, and human fibroblasts. Interestingly, the gene expression pattern from generated neurons most closely resembles that of hPSC-derived peripheral neurons and brain cholinergic neurons. On the other hand, enteric neurons, fibroblasts, epithelial cells, and neural crest sensory neurons remained quite distinct from the generated airway neurons with the current protocol (Figure 3K).

Next, we investigated whether the generated neurons were functional using a multi-electrode array (MEA). Recordings of a MEA enable the measurement of neurons’ spontaneous firing to validate neuronal maturation. Over the course of day 25 through day 73, neurons became more spontaneously active (Figure 3L). Together, the above results indicate that the generated airway cholinergic neurons are mature and functional, including cholinergic muscarinic and nicotinic receptors.

### hPSC-derived airway cholinergic neurons can be co-cultured with ASM

To bring the generated neurons closer to a more biologically and functional relevant model, we investigated whether the generated neurons could be cultured in the presence of cells from the local micro-environment. ASM cells were chosen for co-cultures because smooth muscle-neuronal interactions drive neuronal development and innervation *in utero*^12^. The airway cholinergic neurons were cultured together with immortalized ASM cells from day 35 on (Figure 4A-B) in the same medium used for the neuronal differentiation. IF staining of β-3-tubulin and phalloidin captured neuronal innervation of ASM cells by airway cholinergic neurons (Figure 4C). The formation of VAChT^+^ cholinergic neurons was also confirmed in co-culture with ASM cells (Figure 4D). Additionally, synapse formation of the neurons was observed (Figure 4E). When determining the number of β-3-tubulin^+^-ChAT^+^ cells, it even appeared that co-culturing tends to enhance neuronal maturation: mono-cultures of generated neuron show a proportion of 27.6% (SEM ± 4.5%) β-3-tubulin^+^-ChAT^+^ cells, compared to 40.8% (SEM ± 9.3%) for co-cultured neurons (Figure 4F).

**Figure 4.**
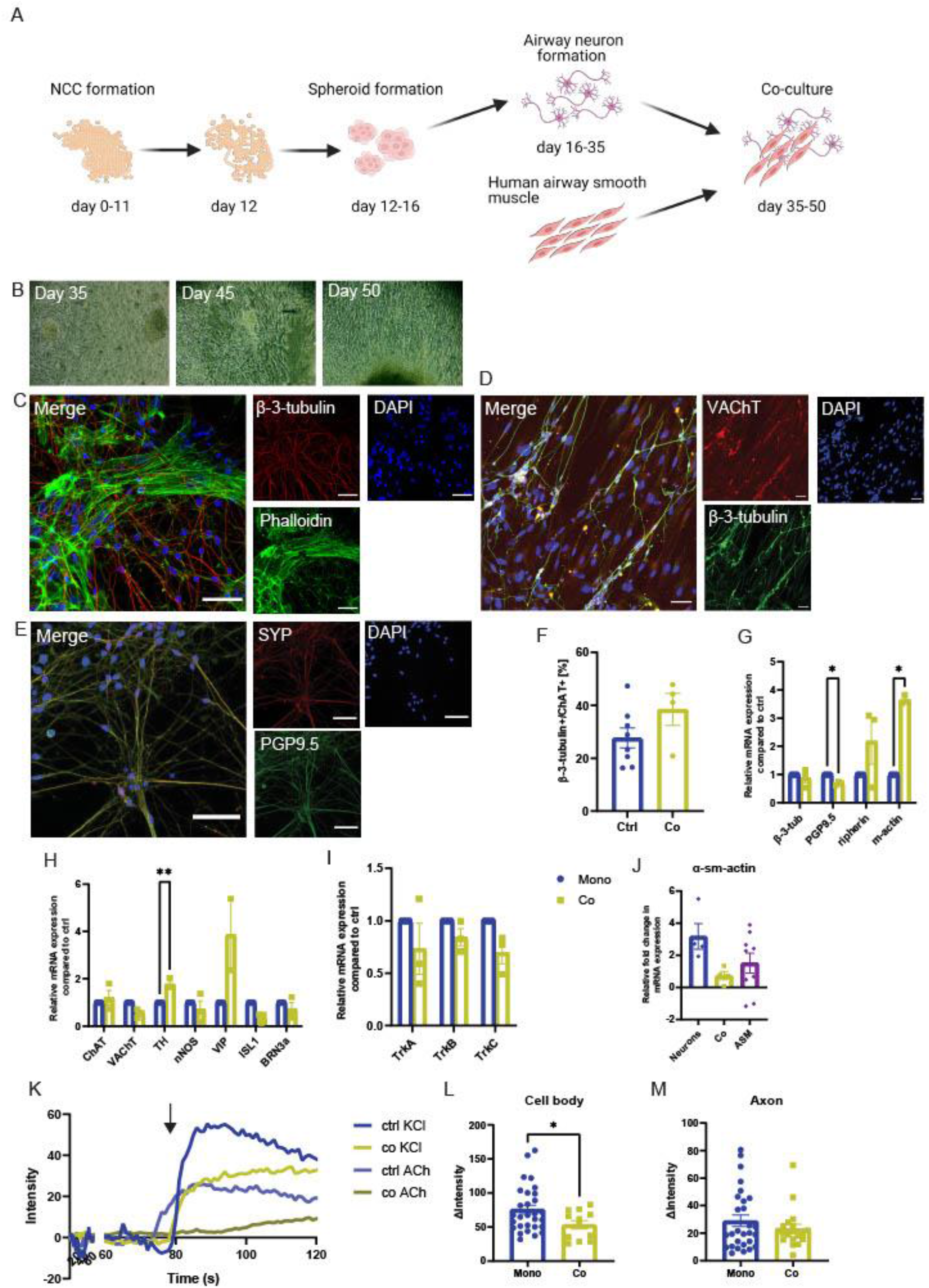
Co-cultures of airway cholinergic neurons with ASM cells. A. Schematic overview of hPSC differentiation into airway cholinergic neurons in the presence of ASM. After 35 days of differentiation, ASM and neurons are cultured together up to day 50. Created with BioRender.com. B. Brightfield images of co-cultures of neurons and ASM over the course of day 35-45-50. C-E. Immunofluorescence images characterizing neurons after co-cultures. β-3-tubulin or PGP9.5 were used as a pan-neuronal marker. The neurons appear well innervated with the ASM (C). ASM is revealed by phalloidin. In addition, also in co-cultures neurons show the presence of VAChT (D) and SYP (E) after 50 days of differentiation. F. FACS analysis demonstrated that for monocultures 27.7% (± 3.8%) of the cells are β-3-tubulin^+^-ChAT^+^, whereas co-cultures are 38.5% (± 6.0%) of cells are β-3-tubulin^+^-ChAT^+^. (N = 4-8) G-I. Gene expression of neurons showed a increased expression of α-sm-actin, and a decrease in PGP9.5 expression (G). Furthermore, TH expression was increased whereas VAChT expression was decreased (H). (N = 4) J. α-sm-actin compared in co-cultures and ASM (N = 4-9) K-M. Live-cell imaging of neurons showed that both in mono- and in co-cultures cells are responsive to KCl and ACh. (K). L. Neuronal cell bodies in both mono-and co-cultures respond to KCl, co-culturing diminishes the response to KCl. (N = 2-6, n = 15-29) M. Axons of hPSC-derived airway neurons show a similar Ca^2+^ response both in mono- and in co-cultures. (N = 2-6, n = 16-28) Figures were presented as mean ± SEM. A paired or unpaired t-test was performed to calculate statistical significance of differences between means for two variables, and a mixed model with a Dunn’s multiple comparison test for three variables; *p<0.05, **p<0.01.

Similar to the characterization of airway cholinergic neuron mono-cultures, we analyzed gene expression in ASM-neuronal co-cultures. As expected, an increase in α-sm-actin mRNA expression was observed, in combination with a decrease in *UCHL1*, since a proportion of the RNA belongs to ASM cells (Figure 4G). α-sm-actin (*ACTA2*) expression in co-cultures did not differ from ASM monocultures (Figure 4J). However, a trend towards more *ACTA2* expression is present. Gene expression of *TUBB3, PRPH*, and *CHAT* did not differ between the conditions. Also, other autonomic neuronal markers, like *NOS1, VIP*, and sensory neuronal markers like *ISL1* and *POU4F1*, did not significantly differ between the two groups (Figure 4H). Only *TH* showed a higher expression in co-cultures, and a clear trend for β-3-tubulin^+^-ChAT^+^ cells (Figure 4H). Trk receptors showed no differences in gene expression (Figure 4I).

Mature neurons show a calcium (Ca^2+^) response when chemically stimulated. Ca^2+^ is a universal second messenger that regulates the most important activities of all cells. It is of critical importance for neuronal activity and neurotransmitter release as it participates in transmitting the depolarizing signal and contributes to synaptic activity and plasticity. Live cell Ca^2+^ imaging using KCl and ACh as stimuli showed clear Ca^2+^ signals in the generated ANs after 50 days of differentiation (Figure 4K). Figure 4K depicts a typical trace after electrochemical stimulation, showing a fast acute transient increase in intracellular Ca^2+^. We calculated Ca^2+^ traces for both the cell bodies and axons: a higher relative intensity peak corresponds to a more intense increase in intracellular Ca^2+^. KCl shows a higher Ca^2+^ response compared to ACh (Figure 4K).

We further examined the intracellular Ca^2+^ signaling response for mono- and co-cultures when stimulated with KCl (Figure 4L-M). Despite the higher number of cholinergic neurons, differentiating neurons in the presence of ASM cells did not result in higher intracellular Ca^2+^ signals when analyzed in the neuronal cell bodies. Neuronal cells showed a relative intensity peak of 75.7 (SEM ± 6.3) in response to KCl, whereas neurons co-cultured with ASM cells showed a relative intensity peak of 53.2 (SEM ± 6.08) (Figure 4L). Interestingly, when comparing the relative intensity peaks of the axons, no difference was observed (Figure 4M); neurons showed a relative Ca^2+^ intensity peak of 29.0 (SEM ± 4.1) in response to KCl, compared to 23.2 (SEM ± 3.3) for co-cultures. No differences in electrochemical responses were observed between mono- and co-cultures upon ACh stimulation, neither for cell bodies nor for axons (Supplementary Figure 4A-B). Co-culturing of the obtained neurons is thus possible with ASM cells and even seems to enhance their differentiation, and the presence of ASM cells changes their Ca^2+^ signaling properties.

### IL-4, IL-13, and IL-33 do not induce neuroplasticity in hPSC-derived airway cholinergic neurons

Neuroplasticity, characterized by an excessive neuronal density and increased firing, is a known phenomenon in asthma^2^. To determine if neuroplasticity can be mimicked *in vitro* using inflammatory cytokines known to be increased in asthma, we stimulated the neuronal cultures with a mix of IL-4, IL-13, and IL-33 during the final five days of differentiation. (Figure 5A). Application of the cytokine mix did not alter the morphology of the neuronal networks (Figure 5B-C). Gene expression analysis confirmed that the profile of genes related to the neuronal network was not affected by the cytokine mix treatment. Neuronal markers did not show a difference in gene expression of several neuronal markers (Figure 5D-F), only TrkA was significantly upregulated. Neither did the number of formed β-3-tubulin^+^-ChAT^+^ cells differ between control and cytokine-stimulated cultures (Figure 5G).

**Figure 5.**
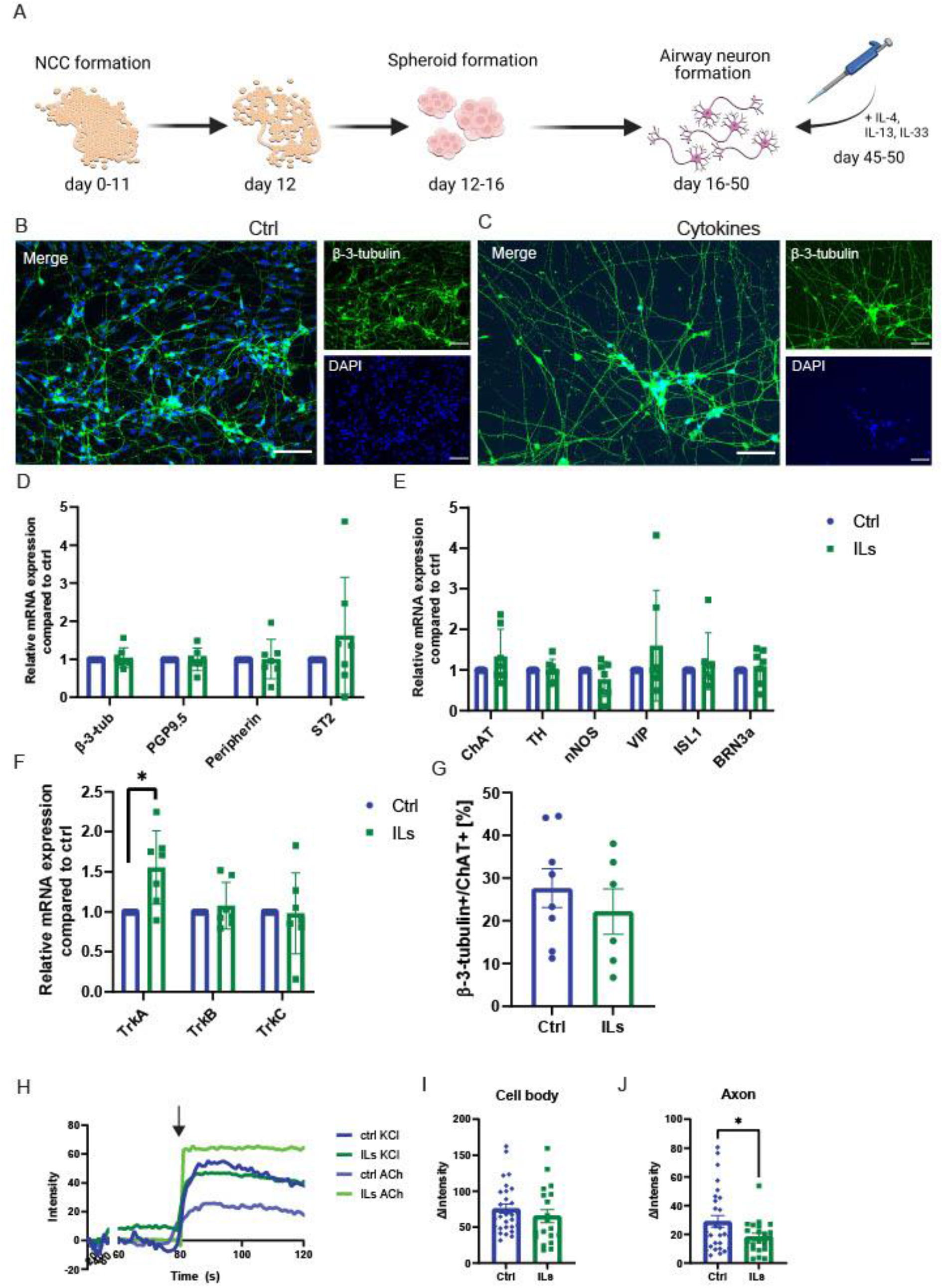
Cultures of cytokine-stimulated airway cholinergic neurons. A. Schematic overview of hPSC differentiation into airway cholinergic neurons with additional cytokine stimulation (IL-4, IL-13, IL-33). Cytokines were added once to the medium at day 45 of differentiation during the final medium change. Created with BioRender.com. B-C. Immunofluorescent staining of β-3-tubulin providing an overview of neuronal-network formation of ctrl neurons compared to cytokine-stimulated neurons. Upon stimulation the cholinergic neurons did not show a different phenotype compared to ctrl neurons. D-F. Gene expression of neurons showed that cytokine stimulation does not alter the expression of neuronal genes in hPSC-derived airway cholinergic neurons. G. FACS analysis demonstrated that for monocultures 22.2% (± 5.3%) of the cells are β-3-tubulin^+^-ChAT^+^, whereas cytokine-stimulation resulted in 20.9% (± 2.3%) of cells as β-3-tubulin^+^-ChAT^+^. H-K. Live-cell imaging of neurons showed that both ctrl and cytokine-stimulated cultures are responsive to KCl. H-I. Neuronal cell bodies in both ctrl and stimulated cultures respond to KCl. (N = 4-5, n = 19-29) J-K. Axons of hPSC-derived airway neurons showed a diminished Ca^2+^ response after cytokine stimulation. (N = 4-5, n = 22-28) Figures were presented as mean ± SEM. A paired or unpaired t-test was performed to calculate statistical significance of differences between means; *p<0.05, **p<0.01.

Moreover, live-cell Ca^2+^ imaging (Figure 5H) did not show a difference in intracellular Ca^2+^ signals measured in the neuronal cell bodies after cytokine stimulation compared to control conditions (Figure 5I), while the Ca^2+^ signals in the axons were significantly lower (*p* < 0.05, Figure 5J). Scanning the appropriate cytokine receptors revealed that only the receptors for IL-4 (*IL4R*) and IL-13 (*IL13RA1*) were present, whereas the IL-33 receptor (*IL1RL1*) was not (Supplementary Table 1). Altogether, it appeared that inflammatory cytokines could not evoke neuroplasticity *in vitro* in the obtained neurons.

### hPSC-derived airway cholinergic neurons show less neuronal activity in response to dexamethasone

Glucocorticosteroids, an established asthma therapy known as inhaled corticosteroids (ICS), reduce airway inflammation and improve lung function. Interestingly, the glucocorticoid receptor (GR) *NR3C1* was detected in differentiated and mature neurons, implying a potential response to glucocorticosteroid stimulation (Supplementary Table 1). We used dexamethasone, a powerful corticosteroid, to gain more insight into the effect of glucocorticosteroids on neuroplasticity in airway cholinergic neurons. We stimulated the generated neurons with dexamethasone during the final five days of differentiation (Figure 6A). Both the control and the dexamethasone-stimulated neuronal networks appeared normal and healthy (Figure 6B-C). A gene expression analysis of several neuronal markers showed that dexamethasone-stimulation did not alter the expression of the differentiated neurons (Figure 6D-F). Besides, the number of β-3-tubulin^+^-ChAT^+^ neurons formed did not differ significantly: 27.6% (SEM ± 4.5%) β-3-tubulin^+^-ChAT^+^ cells, compared 19.7 % (SEM ± 3.7%) for stimulated cultures. However, a trend towards fewer neurons formed was observed (Figure 6G).

**Figure 6.**
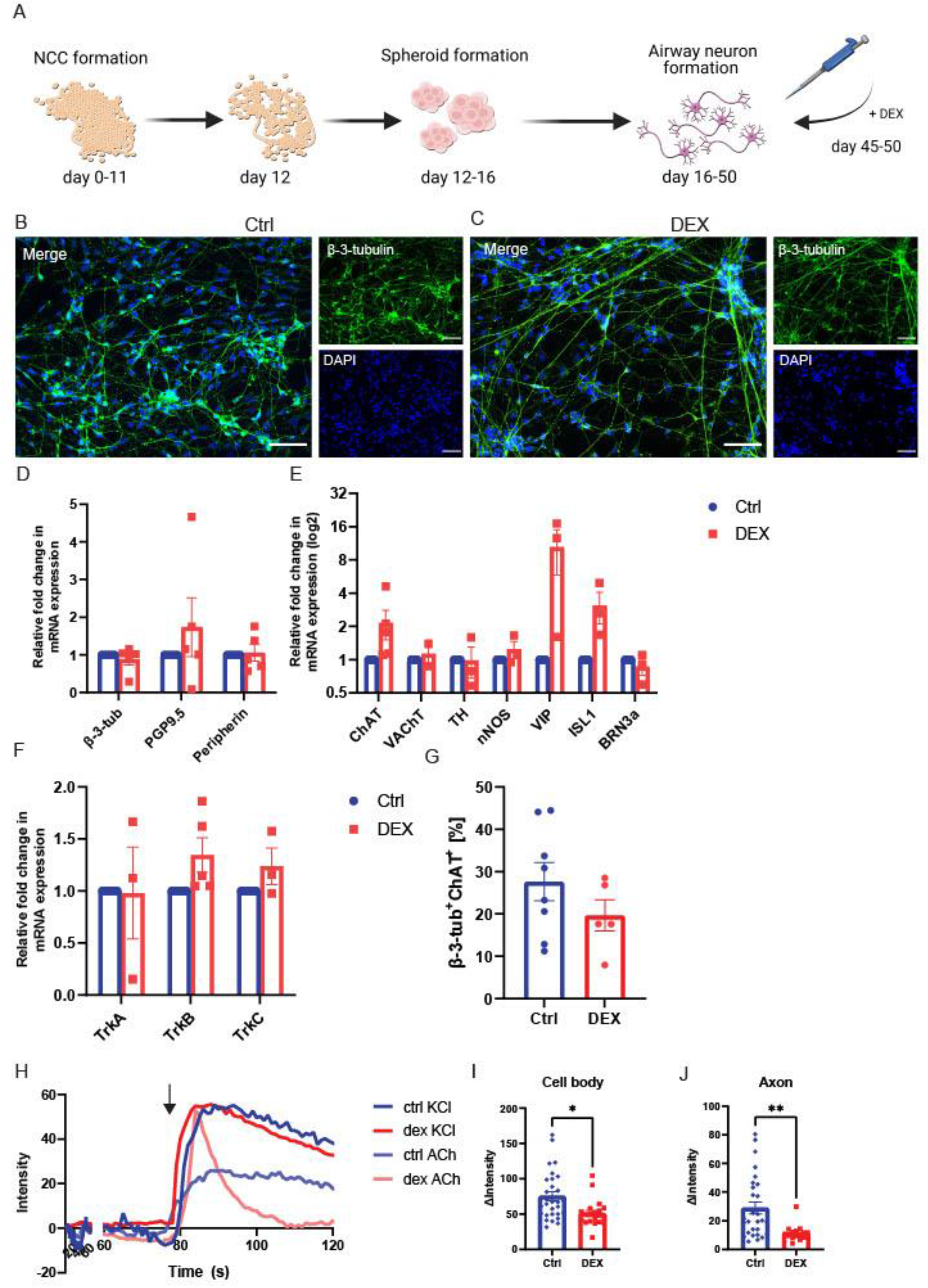
Cultures of dexamethasone-stimulated airway cholinergic neurons. A. Schematic overview of hPSC differentiation into airway cholinergic neurons with additional dexamethasone stimulation. Dexamethasone was added once to the medium at day 45 of differentiation during the final medium change. Created with BioRender.com. B-C. Immunofluorescent staining of β-3-tubulin providing an overview of neuronal-network formation of ctrl neurons compared to dexamethasone-stimulated neurons. Upon dexamethasone-stimulation the cholinergic neurons do not show a different phenotype compared to ctrl neurons. D-F. Gene expression of neurons showed that dexamethasone stimulation does not alter the expression of neuronal genes in hPSC-derived airway cholinergic neurons. G. FACS analysis demonstrated that for monocultures 27.7% (± 3.8%) of the cells are β-3-tubulin^+^-ChAT^+^, whereas dexamethasone-stimulation resulted in 20.9% (± 2.3%) of cells as β-3-tubulin^+^-ChAT^+^, with a tendency to decrease cholinergic neurons. H-K. Live-cell imaging of neurons showed that both ctrl and dexamethasone-stimulated cultures are responsive to KCl. H-I. Neuronal cell bodies in both ctrl and dexamethasone-stimulated cultures respond to KCl, in dexamethasone-stimulated cultures the response to KCl diminished. (N = 4-5, n = 20-29) J-K. Axons of hPSC-derived airway neurons showed a diminished Ca^2+^ response after dexamethasone stimulation. (N = 4-5, n = 15-28) Figures were presented as mean ± SEM. A paired or unpaired t-test was performed to calculate statistical significance of differences between means; *p<0.05, **p<0.01.

Despite the number and the type of differentiated cells seemingly unchanged, live-cell Ca^2+^ imaging showed a lower response to KCl in dexamethasone-stimulated neurons than control conditions (Figure 6H-J). Following the Ca^2+^ traces, the dexamethasone-stimulated cells responded to KCl stimulation (Figure 6H), albeit with lower detected intracellular Ca^2+^ levels. The Ca^2+^ signals in cell bodies dropped from a 75.7 (SEM ± 6.3) intensity peak to 51.2 (SEM ± 4.3) in dexamethasone-stimulated cultures (*p* < 0.05, Figure 6I), and the Ca^2+^ response in axons dropped from a peak of 29.0 (SEM ± 4.1) intensity peak to 11.5 (SEM ± 1.5) (*p* < 0.01, Figure 6J). Interestingly, Ca^2+^ signals in cell bodies increased in response to ACh after dexamethasone stimulation ((*p* < 0.01, Supplementary Figure 4C), but not in axons (Supplementary Figure 4D). Together, this indicated that the generated neurons are responsive to the effects of the corticosteroid dexamethasone.

## Discussion

This study demonstrated the differentiation of airway cholinergic neurons via a vagal NCC precursor. The cholinergic phenotype, mainly present in the airways, was confirmed in our cultures by ChAT and VAChT expression. Predominant *NTRK2* expression further confirmed the airway neuronal phenotype. Next, we improved the translational value of the model by co-culturing differentiated airway cholinergic neurons in the presence of ASM cells. Moreover, the generated neurons showed successful implementation as a drug-screening tool as dexamethasone stimulation diminished neuronal excitability capabilities.

The induction of the vagal precursor was highly efficient, with a >90% yield of HNK^+^-p75^+^ cells. This high efficiency is in accordance with vagal NCC induction for enteric neuronal development, showing a similar yield for NCC induction^17^. The subsequent differentiation of vagal NCC into airway cholinergic neurons showed the formation of a neuronal network that kept increasing gradually, reflected both in protein and in gene expression. Protein expression analysis revealed that most neurons displayed a cholinergic phenotype. IF staining showed the utmost part of neurons to be VAChT^+^, while only a minor part was TH^+^. In addition, FACS analysis revealed ChAT expression in the majority of β-3-tubulin^+^ cells. Comparing the generated neurons with other cells using a PCA affirmed that the characterization mainly overlaps with peripheral and brain cholinergic neuronal samples. Similarly, RNAseq analysis showed that neurons were the predominant cell type in our samples. VAChT expression supported maturation of the airway neurons over time: at day 35, VAChT is hardly present, while at day 50, VAChT is present in the majority of neurons. An intracellular Ca^2+^ response to electrochemical stimulus further confirmed neuronal maturation.

Electrophysiological patch-clamp recordings have demonstrated neuronal firing of hPSC-derived peripheral sensory neurons (SN). 90% of cells of a differentiated SN batch fired at least one action potential (AP)^35^. Likewise, hPSC-derived sympathetic neurons possess electrophysiological properties similar to rodent sympathetic neurons, as shown in phasic/tonic APs^36^. While the hPSC-derived airway neurons showed spontaneous activity after 50 days of differentiation on a MEA system, it has been shown before that hPSC-derived peripheral SNs showed spontaneous neural activity at day 25, with a peak in burst and firing activity at day 65, measured by a similar MEA system^37^. Besides, the hPSC-derived peripheral SNs did not synchronize network activity^37^. We obtained similar results with the spontaneous firing of the generated neurons from day 25 until day 73, with increasing activity over time and without synchronized firing.

Following the changes in gene expression, the neuronal differentiation follows a developmental maturation as observed *in vivo*. TH is considered a marker for sympathetic neurons. However, it is also a precursor for airway cholinergic neurons during development^27^. TH is expressed transiently during development: in neurons and neuroendocrine cells, TH is expressed in cells that in adulthood no longer express TH or only at very low levels^27^. The peak expression of *TH* observed at day 35 of differentiation could be explained by a transient TH expression. The obtained neurons still express *TH* after 50 days of differentiation, directing towards an incomplete neuronal maturation. However, IF staining showed a minor proportion of cells were TH^+^, which could be the transient expression or a small population of sympathetic neurons.

To validate the generated neurons as a human *in vitro* model using a reference drug, we assessed the response to corticosteroids. As the GR *NR3C1* is present, we expected the potent glucocorticosteroid dexamethasone to exert an effect on the neurons. Dexamethasone was shown before to impair brain development in murine models, whilst not affecting other somatic structures^38^. Also, dexamethasone stimulation of PC12 cells resulted in reduced neurite formation, and also decreased cholinergic phenotype by suppression of acetylcholine synthesis^39^. In accordance with impairment of neuronal development, in the generated neurons, the cellular response to KCl during live-cell Ca^2+^ imaging was clearly diminished both in the cell bodies and the axons. The difference in cholinergic neuronal yield and network formation did not seem affected. Currently, much is unknown about the effects of glucocorticosteroids on neuronal development and plasticity, as their effects have mainly been studied in relation to inflammation. Recently, the effects of glucocorticosteroids on neuronal development have become more interesting; as for PC12 cells, many binding GR sites are available^40^. However, this is a rat cell line, and provides limited information on the human situation, especially for the respiratory tract. The generated airway cholinergic neurons can serve as a human *in vitro* disease model to understand more about the effects of glucocorticosteroids on airway neuronal plasticity.

The presence of multiple cell types gives a more accurate response to stimulation of any kind if the model is applied as a disease model. Co-culturing the generated neurons with cells of the local microenvironment improved the model’s value, as cellular interaction can be incorporated. We found that the hPSC-derived airway neurons can be co-cultured with ASM cells without any difficulties. The neurons innervated the ASM cells, and the percentage of β-3-tubulin^+^-ChAT^+^ cells was higher in co-cultures than in mono-cultures. Co-cultures of obtained peripheral neurons with intestinal cells or cardiac cells lead to accelerated maturation by showing a greater action potential at an earlier time point during differentiation^16,36^. However, the generated neurons did not display an enhanced response to KCl stimulation.

In the generation of a model for neuroplasticity in asthma, the presence of neurons expressing a subset of nicotinic and muscarinic receptors that respond to ACh is essential. Moreover, interactions with cells like ASM or mast cells are important. ASM-neuronal interactions seem to rely on BDNF and TrkB^8^, but many of the underlying mechanisms are currently unknown. We previously demonstrated that IL-33 can trigger neuronal plasticity in co-cultures of ASM cells and neurons, but not in mono-cultures of either of the two cell types^41^. We did not observe profound effects of cytokine stimulation to induce neuronal plasticity following inflammation. Interactions with other cell types might be important for the induction of neuronal plasticity, that can simply not be mimicked in neuronal cultures alone. This finding might explain that we could not induce augmented network formation here but that the presence of ASM or other cells is essential. The addition Th2 cytokines to ASM-neuronal co-cultures might be a logical next step, as well as the addiction of mast cells to the generated neurons to study neuro-immune interactions *in vitro* as inflammation is often underlying neuroplasticity^2^. This information is valuable for the study of neuronal plasticity in asthma and eventually using multi-cell-culture models for better predictive *in vitro* models.

In conclusion, we have demonstrated the ability to differentiate hPSCs into airway cholinergic neurons via a vagal precursor using chemically defined media. The protocol for airway cholinergic neurons is a valuable addition to the protocols for sympathetic, enteric, and sensory neurons developed already. The neurons provide a helpful platform for the study of neuronal plasticity, which can be expanded by co-culturing with cells of the local microenvironment. Furthermore, the neurons have proven to be used as a disease model and a platform for drug studies. These hPSC-derived airway neurons help identify mechanisms underlying neuronal plasticity and potentially contribute to personalized medicine.

## Materials and methods

### H9WA09 cell culturing

H9WA09 cells were obtained from the European Institute for the Biology of Ageing (ERIBA) at the University of Groningen. H9WA09 cells were cultured on Matrigel hESC-qualified (Corning, 354277) pre-coated 6-well plates (6WP) in mTeSR1 medium (STEMCELL technologies, 85850). The cells were incubated at 5% CO2 and 37°C. Once the cells grew confluent, the H9WA09 cells were passaged using ReLeSR (STEMCELL technologies, 05872). The pluripotency of the H9WA09 cells was tested regularly by staining for the pluripotency-marker OCT4. In addition, the cells were examined regularly for the presence of mycoplasma.

### Differentiating H9WA09 cells towards a neuronal cell fate

For differentiation of H9WA09 cells into airway cholinergic neurons, several stages were passed (Supplementary Table 2). Vagal NCC induction was started when pluripotent stem cells were 40-60% confluent. Pluripotent stem cells were first differentiated into vagal NCCs in 12 days. Two types of media were used: KSR medium (KnockOut DMEM (Thermo Fisher, 10829018) and 15% KnockOut Serum Replacement (Thermo Fisher, 10828028)) and N2 medium (DMEM HEPES (Thermo Fisher, 12320032), 1% Penicillin-Streptomycin (Thermo Fisher, 15070063), and 10 µg/ml N2 supplement (Thermo Fisher, A1370701)). The manufacturer supplies N2 supplement as 100x, but double the amount was used in this protocol. KSR and N2 medium were freshly supplemented with 10 µM SB431542 (STEMCELL Technologies, Vancouver, Canada, 72234), 1 µM LDN193189 (STEMCELL Technologies, Vancouver, Canada, 72147), 3 µM CHIR99021 (STEMCELL Technologies, Vancouver, Canada, 72054), and 1 µM retinoic acid. The medium was changed every other day, medium composition according to Supplementary Table 3.

Next, the vagal NCCs were cultured in the form of floating spheroids for four days. The cells were washed with EDTA (0.5 mM) twice, followed by 10 min incubation, 37°C. After aspirating the EDTA, vagal precursor (VP) spheroid medium was added, consisting of Neurobasal Medium (Thermo Fisher, 21103049) supplemented with 10 µl/ml N2 supplement, 20 µl/ml B27 supplement (Thermo Fisher, 17504044), 10 µl/ml Glutamax (Thermo Fisher, 35050061), 10 µl/ml MEM Nonessential Amino Acids). 10 ng/ml FGF2 (Thermo Fisher, PHG6015), and 3 µM CHIR99021 (STEMCELL Technologies, 72054), was freshly added. The cells were mechanically detached in VP spheroid medium using a serological pipette before transfer to a 6WP pre-coated with anti-adherence solution (STEMCELL Technologies, 07010). Detached cells from five wells of a 6WP were added divided over six wells of a 6WP. On day 14, the medium was refreshed.

After spheroid formation, the vagal cells undergo airway cholinergic neuron induction and airway cholinergic neuron maturation. On day 16 of the protocol, the VP spheroid medium was aspirated, and the spheroids were dissociated using EDTA (0.5 mM; wash twice, followed by 10 min incubation, 37°C). The cell suspension was carefully transferred to a tube and centrifuged (290 g, 1 min, RT). The supernatant was aspirated, and the cell pellet was resuspended in airway neuron (AN) medium (Neurobasal medium supplemented with, 10 µg/ml N2 supplement, 20 µl/ml B27 supplement, 10 µg/ml Glutamax, and 10 µg/ml MEM Nonessential Amino Acids). 10 ng/ml BDNF (Peprotech, 450-02) and 100 µM freshly prepared L-Ascorbic Acid (Sigma-Aldrich, A5960) were added before medium change. Dissociated spheroids were plated onto culture plates pre-coated with 15 µg/mL Poly-L-Ornithine (PLO, Sigma-Aldrich, P4538), 2 µg/mL fibronectin (FB, Thermo Fisher, 33016015), and 2 µg/mL laminin (LM, R&D systems, 3400-010-02).

In the first stage of airway cholinergic neuron induction (day 16-30), the medium was changed three times per week, changing ¾ of the total volume. In the second stage (day 30-40), the medium was changed twice per week the medium volume was 1.5 times increased. In addition, from day 35 onwards, AN medium was supplemented with 2 µg/ml FB and 2 µg/ml LM. In the final stage of airway cholinergic neuron maturation (day 40-50), the medium was changed once per week. Cells were stimulated with 10 nM dexamethasone, or with the cytokines IL-4 (10 ng/mL, Peprotech, 200-04), IL-13 (3 ng/mL, Peprotech, 200-13), and IL-33 (10ng/mL, Peprotech, 200-33). PCR analysis and immunofluorescence (IF) staining were performed on different time points: day 25, 35, and day 50. Additionally, fluorescence-activated cell sorting (FACS) was performed on day 50 (Figure 2).

### ASM culture

Immortalized primary human ASM cells were cultured in DMEM supplemented with 10% Fetal Bovine Serum (FBS), 2,2% Penicillin-Streptomycin, and 0,6% Amphotericin B (Thermo Fisher, 15290026). Once the cells grew 100% confluent, the ASM cells were passaged using 0.25% trypsin (Thermo Fisher, 25200056). The cells were incubated at 5% CO2 and 37°C. The cells were tested frequently for the presence of mycoplasma.

### Co-culturing hPSCs and ASM

hPSC-derived airway cholinergic neurons were cultured together with ASM cells from day 35 of differentiation on to generate co-cultures. ASM cells were treated with 40 µg/ml mitomycin C (Thermo Fisher, J63193-MA) to attenuate proliferation, 2 hours at 37°C. After 2 hours, the ASM cells were cultured in starvation medium (DMEM + 1% Insulin-Transferrin-Selenium (ITS, Thermo Fisher, 41400045) + 2,2% Penicillin-Streptomycin + 0,6% Amphotericin B) for at least one hour before use. ASM cells were plated (50.000 cells/cm^2^) in pre-coated PLO/FB/LM plates and cultured overnight. On day 35, the hPSC-derived neurons were dissociated using EDTA (0.5 mM; wash twice, followed by 10 min incubation, 37°C) and added to the ASM. The hPSC-derived neurons of one well were replated in a 1:1 ratio in a new well.

### Gene expression hPSC-derived neurons

mRNA was isolated using NucleoSpin RNA XS kit (Macherey-Nagel, 740902.50) according to the manufacturer’s protocol. The yield and purity of the isolated RNA were measured using the NanoDrop 1000 spectrophotometer. The RNA was either processed for RT-qPCR or sent for RNA sequencing (RNAseq).

For RT-qPCR, cDNA was synthesized using Reverse Transcription System (Promega, A3500) according to the manufacturer’s protocol. The RTq-PCR reactions were completed using SYBR Green. A list of the qPCR forward primers and reverse primers used is provided in Supplementary Table 4. The program for RT-qPCR reactions started with polymerase activation at 95°C for 10 minutes, 45 cycles of PCR cycling, which included denaturation at 95°C for 30 seconds, annealing for 30 seconds at 59°C, and extension at 72°C for 30 seconds, and incubation at 72°C for 5 minutes. Melting curves were obtained consecutively: 15 seconds at 95°C, 15 seconds at 55°C, and 15 seconds at 95°C. Analysis of the gene expression was performed with Quantstudio Real-Time PCR software v1.2^19^.

RNA from generated neurons was sent to GenomeScan BV (Leiden, Netherlands) for sequencing, and raw counts were provided together with a quality report. Genes with at least 1 count per million (CPM) were considered expressed and included for further analysis. Non-expressed genes were removed. The gene level read counts were normalized using DEseq2 in R (version 4.1.0)^20^, and normalized expression levels of several neuronal and non-neuronal markers was compared. Normalized counts can be found in Supplementary file 1. Principal component analysis (PCA) analysis was performed using the R packages: “edgeR” and “limma” and plotted using “ggplot”, creating multidimensional scaling plots.

### Immunofluorescence staining of cultures

Cells were fixed in 4% paraformaldehyde (PFA, Sigma-Aldrich, 97H0752) and permeabilized, using 0,3% Triton X (Sigma-Aldrich, 101371900) for 5 minutes, RT. The cells were blocked for 1 hour with blocking buffer (Cyto-TBS + 2% bovine serum albumin (BSA, Sigma Aldrich, 1002695029)) and incubated with a primary antibody overnight at 4°C. See Supplementary Table 5**Error! Reference source not found**. for used antibodies and dilutions. The next day, the cells were incubated with a secondary antibody for 2 hours in the dark, RT. Optionally, cells were counterstained for 45 minutes using 1 unit/assay Alexa Fluor 488 Phalloidin (Thermo Fisher, A12379). Cells were mounted using a mounting medium with DAPI (Abcam, ab104139). Samples were imaged using a Leica DM4000b (Leica microsystems) or a Zeiss LSM 780 (Zeiss, Germany) microscope and analyzed using Fiji (http://fiji.sc/).

### Fluorescence-Activated Cell Sorting

Fluorescence-Activated Cell Sorting (FACS) was performed on day 12 and day 50 of the differentiation protocol. On day 12, FACS was performed on live cells to determine the percentage of p75^+^-HNK1^+^ cells. Samples were washed with PBS and incubated with Accutase (Sigma Aldrich, A6964) for 15 minutes at 37°C. The cells were resuspended in Accutase and centrifuged for 5 minutes at 200 x g. Subsequently, the cells were washed with FACS buffer (PBS supplemented with 0.5% BSA and 20 mM Glucose) and then centrifuged for 5 minutes at 200 x g. Cells were incubated using HNK1 (Biolegend, 359613) and p75 (Biolegend, 345101) for 60 minutes on ice. After incubation the cells were centrifuged at 200 x g for 5 minutes and resuspended in FACS Buffer. 7-aminoactinomycin D (7-AAD, Thermo Fisher, A 1310) was added 1:50 as a cell death marker.

On day 50, FACS was performed on fixed cells to determine the percentage of β-3-tubilin^+^-Choline acetyltransferase (ChAT)^+^ cells. Samples were washed twice with EDTA (0.5 mM) and incubated with EDTA (15min, 37°C). The cells were mechanically transferred using a serological pipette into FACS tubes and centrifuged (290 g, 1 min, RT). A Fix & Perm™ Cell Permeabilization kit (ITK diagnostics, GAS-002) was used according to the manufacturer to fixate and permeabilize the cells. PE-anti-β-3-tubulin (Biotechne, NB600-1018PE) and APC-anti-Choline Acetyltransferase (Abcam, ab224001) antibodies were incubated for 40 min. Samples were then analyzed using the BD LSR II system (BD Biosciences). Supplementary Table 5 shows the antibodies and the dilutions that were used. Data was acquired using Diva 8.0 Software.

### Live cell Ca^2+^ imaging using Fluo-4-AM

Live-cell imaging to show mature neuronal response to potassium chloride (KCl) or acetylcholine (ACh) was performed using Fluo-4-AM (Invitrogen™, F14217). Cells were incubated with Fluo-4-AM in HBSS-Ca^2+^ (45 min, RT, dark) and maintained at RT in the dark until data acquisition. Data were acquired using the Zeiss LSM 780 microscope. 3-10 neurons were identified for data acquisition; the cells should display a typical neuronal morphology with dendritic and axonal processes recognizable by cellular polarity and proportionate size. Cells were excited with 488-nm light (for Fluo-4-AM) and red-nm light (for 7-ADD cell death marker). Images were collected by taking an image every 100 msec for 3 minutes. After 30 seconds, the cells were challenged by adding 60 mM KCl or 100 μM ACh. An increase in intensity was measured to quantify the neuronal response. Supplementary videos are provided to demonstrate the calcium response (Supplementary video 1-8).

### Spontaneous firing of neurons

A Maestro Pro (Axion Biosystems) multi-electrode array (MEA) system was used to measure the spontaneous firing of neurons^21^. We coated CytoView MEA 48 plates (Axion Biosystems, M768-tMEA-48W) containing sixteen embedded electrodes per well with PO/LM/FN. The coating solution was aspirated, and the wells were left to dry for 30 minutes. Dissociated NCC spheroids (day 16) were seeded in high-concentration LM (10 µM/mL) drops (5 µL/drop) into the center of the MEA well. The cells were incubated (60 min, 37°C) before adding AN medium. Repeated recordings were made every 10 days for 15 minutes. The MEA plate was inserted into the MEA Maestro (37C, 5% CO2) for spike detection. Axion AxIS Software recorded raw voltage data and detected spikes for rate analysis.

### Statistical analysis of H9WA09 differentiated cells

The data are presented as mean ± standard error of the mean (SEM). Statistical differences between distinct conditions were calculated using a two-way ANOVA or mixed effect analysis followed by either a Dunnett’s test or a Tukey’s multiple comparisons test to calculate significant differences comparing three or more variables. A paired t-test was performed comparing two variables. Statistical analyses were performed in GraphPad Prism (version 9.3.0), and performed tests are specified in the figure legends.

## Acknowledgments

The work was financially supported by the More Knowledge with Fewer Animals (*Meer Kennis met Minder Dieren*) programme of ZonMW (grant number 114021505) with co-financing from Stichting Proefdiervrij, Aquilo BV, Boehringer Ingelheim and Longfonds. We would like to thank Floris Foijer (ERIBA, Groningen, the Netherlands) for the cells he kindly provided, and Marina Trombetta Lima (University of Groningen, Groningen, the Netherlands) for her technical assistance.

## List of abbreviations

6WP: 6 wells-plate
a.a.: ascorbic acid
ACh: Acetylcholine
AN: Airway neuron
AP: Action potential
ASM: Airway smooth muscle
BDNF: Brain-derived neurotrophic factor
BSA: Bovine serum albumin
Ca^2+^: Calcium
ChAT: Choline acetyltransferase
CPM: Counts per million
CNS: Central nervous system
FACS: Fluorescence-activated cell sorting
FB: Fibronectin
FBS: Fetal Bovine Serum
GDNF: Glial cell-derived neurotrophic factor
hPSC: human pluripotent stem cell
KCl: Potassium chloride
LM: Laminin
ICS: Inhaled corticosteroids
IF: Immunofluorescence
MEA: Multi-electrode array
NCC: Neural crest cell
NTF: neurotrophic factor
nNOS: Neuronal nitric oxide synthase
PCA: Principal component analysis
PFA: Paraformaldehyde
PLO: Poly-L-Ornithine
PNS: Peripheral nervous system
r.a.: retinoic acid
RT: Room temperature
SEM: Standard error of the mean
SN: Sensory neuron
SYP: Synaptophysin
TH: Tyrosine hydroxylase
TrkB: Tropomyosin receptor kinase B
VAChT: Vesicular acetylcholine transporter
VIP: Vasoactive intestinal peptide
VP: Vagal precursor

